# Genome assembly of the Australian black tiger shrimp (*Penaeus monodon*) reveals a fragmented IHHNV EVE sequence

**DOI:** 10.1101/2021.11.11.468259

**Authors:** Roger Huerlimann, Jeff A Cowley, Nicholas M Wade, Yinan Wang, Naga Kasinadhuni, Chon-Kit Kenneth Chan, Jafar Jabbari, Kirby Siemering, Lavinia Gordon, Matthew Tinning, Juan D Montenegro, Gregory E Maes, Melony J Sellars, Greg J Coman, Sean McWilliam, Kyall R Zenger, Mehar S Khatkar, Herman W Raadsma, Dallas Donovan, Gopala Krishna, Dean R Jerry

**Affiliations:** ARC Industrial Transformation Research Hub for Advanced Prawn Breeding, Australia; Centre for Sustainable Tropical Fisheries and Aquaculture, College of Science and Engineering, James Cook University, Townsville, QLD 4811, Australia; Centre for Tropical Bioinformatics and Molecular Biology, James Cook University, Townsville, QLD 4811, Australia; CSIRO Agriculture and Food, 306 Carmody Road, St Lucia, QLD, 4067, Australia; Australian Genome Research Facility Ltd, Level 13, Victorian Comprehensive Cancer Centre, 305 Grattan St, Melbourne VIC 3000, Australia; Laboratory of Biodiversity and Evolutionary Genomics, KU Leuven, Leuven, 3000, Belgium; Center for Human Genetics, UZ Leuven- Genomics Core, KU Leuven, Leuven, 3000, Belgium; CSIRO Agriculture and Food, Integrated Sustainable Aquaculture Program, 144 North Street, Woorim, QLD 4507, Australia; Sydney School of Veterinary Science, Faculty of Science, The University of Sydney, Camden, NSW 2570, Australia; Seafarms Group Ltd, Level 11 225 St Georges Terrace, Perth, WA 6000, Australia

**Keywords:** *Penaeus monodon*, Australia, genome assembly, PacBio, IHHNV EVE

## Abstract

Shrimp are a valuable aquaculture species globally; however, disease remains a major hindrance to shrimp aquaculture sustainability and growth. Mechanisms mediated by endogenous viral elements (EVEs) have been proposed as a means by which shrimp that encounter a new virus start to accommodate rather than succumb to infection over time. However, evidence on the nature of such EVEs and how they mediate viral accommodation is limited. More extensive genomic data on Penaeid shrimp from different geographical locations should assist in exposing the diversity of EVEs. In this context, reported here is a PacBio Sequel-based draft genome assembly of an Australian black tiger shrimp (*Penaeus monodon*) inbred for one generation. The 1.89 Gbp draft genome is comprised of 31,922 scaffolds (N50: 496,398 bp) covering 85.9% of the projected genome size. The genome repeat content (61.8% with 30% representing simple sequence repeats) is almost the highest identified for any species. The functional annotation identified 35,517 gene models, of which 25,809 were protein-coding and 17,158 were annotated using interproscan. Scaffold scanning for specific EVEs identified an element comprised of a 9,045 bp stretch of repeated, inverted and jumbled genome fragments of Infectious hypodermal and hematopoietic necrosis virus (IHHNV) bounded by a repeated 591/590 bp host sequence. As only near complete linear ~4 kb IHHNV genomes have been found integrated in the genome of *P. monodon* previously, its discovery has implications regarding the validity of PCR tests designed to specifically detect such linear EVE types. The existence of joined inverted IHHNV genome fragments also provides a means by which hairpin dsRNAs could be expressed and processed by the shrimp RNA interference (RNAi) machinery.

## INTRODUCTION

Shrimp aquaculture plays a central role in producing high quality protein for human consumption, with global aquaculture production of the two major species, *Penaeus vannamei* and *P. monodon*, reaching close to six million tons in 2018 (FAO 2020). However, diseases, such as those caused by highly pathogenic viruses, are currently a major contributor to unfulfilled production potential (FAO 2020). Therefore, a more advanced understanding of the host defense mechanisms that suppress infection will be critical to finding solutions to viral diseases (Kulkarni et al. 2021; Hauton 2017; Yang et al. 2021).

Initially described in insects, the viral accommodation mechanism has been hypothesized to explain why farmed shrimp highly susceptible to morbidity and mortality proceeding their initial encounter with a new virus tend to become less susceptible over time (Flegel 2020). Viral accommodation is mediated through host-genome integrated endogenous viral elements (EVEs) that can be inherited after integration into the germ line. The expressed EVE-specific double-stranded RNA (dsRNA) is then processed by the host RNA interference (RNAi) pathway, suppressing viral RNA expression levels and therefore infection loads. In the case of RNA viruses, a linear copy viral DNA (cvDNA) or circular copy viral DNA (ccvDNA) can be reverse transcribed by the host (Taengchaiyaphum et al. 2021). These DNA copies of virus RNA can then either autonomously insert into the host genome to become an EVE, or be used directly as a template for dsRNA transcription as an initial step to RNAi-mediated suppression of virus infection (Taengchaiyaphum et al. 2021).

Of the >50,000 known crustacean species, high-quality genome assemblies are only available for a select few taxa, driven primarily by the commercial or unique biological significance of certain species. Genome assemblies provide a reference base for functional transcriptomic studies (Yue and Wang 2017; Chandhini and Rejish Kumar 2019), aid in the positioning of genetic markers used for selective breeding (Houston et al. 2020; Zenger et al. 2017) and provide an important resource for the examination and characterization of genomic regions of commercial or biological interest (Guppy et al. 2020; Hollenbeck and Johnston 2018). However, crustacean genomes have also proved immensely challenging to assemble due to their large (>2 Gbp), highly repetitive (>50%), and highly heterozygous genomes (Yuan et al. 2021a). To some extent, these difficulties have been alleviated by the advent of single-molecule long-read sequencing and improved genome assemblers. Extracting intact high-quality genomic DNA from muscle tissue of crustaceans like shrimp has also proved problematic and exacerbated difficulties in obtaining high-quality data from various NGS platforms (Angthong et al. 2020). Despite these challenges, genome assemblies highly fragmented into more than a million contigs have been reported for the penaeid shrimp species *P. vannamei* (Yu et al. 2015), *P. japonicus* (Yuan et al. 2018), and *P. monodon* (Van Quyen et al. 2020; Yuan et al. 2018). Through applying long-read sequencing and HiC scaffolding, less fragmented high-quality genomes have also been achieved recently for *P. vannamei* (Zhang et al. 2019), *P. monodon* (pseudo-chromosome level) (Uengwetwanit et al. 2020) and *P. japonicus* (Kawato et al. 2021).

Reported here is a high-quality draft genome assembly of a single-generation inbred male *P. monodon* from eastern Australia, a population genetically distinct from others across their South East Asian, Indo-Pacific and East African distribution (Vu et al. 2021). We report and resolve the genomic structure of an EVE of Infectious hypodermal and hematopoietic necrosis virus (IHHNV) comprised of repeated, inverted, and jumbled IHHNV genome fragments. We discuss the disease detection implications of false PCR-positives for infectious IHHNV, and how the EVE might have originated.

## METHODS & MATERIALS

### Shrimp breeding and selection for sequencing

A second-generation (G2) male *Penaeus monodon* that had undergone a single cycle of inbreeding was selected for genomic sequencing. The original wild-caught broodstock were collected from a Queensland east coast location (approximately 17.3°S, 146.0°E) in September 2013. In October 2013, 14 first-generation (G1) families were produced from the brood stock at Seafarm Flying Fish Point hatchery (approximately 17.5°S, 146.1°E). In February 2015, pleopod tissue was sampled from 50 female and 50 male G1 broodstock. These tissues were genotyped (using 2 × 60 SNP panels (Sellars et al. 2014) to identify the parental origin of each broodstock and to select related mating pairs to generate the inbred G2 progeny. In August 2015, groups of 50 juvenile males from 5 inbred G2 families were euthanized to collect muscle tissue from the first abdominal segment for sequencing and the second most anterior pair of pleopods for genotyping. These tissues, as well as the remainder of each shrimp (archived source of tissue for sequencing) were snap frozen under dry ice pellets and stored at −80°C. Each shrimp was then genotyping using the 120-SNP panel (Sellars et al. 2014) and a genome-wide SNP assay based on DArTSeq (Guppy et al. 2020). After ranking the 50 males based on inbreeding coefficient (F) and multi-locus heterozygosity (MLH) data from the 120-SNP panel, the individual (named Nigel) with the highest inbreeding coefficient was chosen for genomic sequencing. The choice was confirmed using a genome-wide SNP assay based on DArTSeq of the top five inbred shrimp based on the 120-SNP panel which recovered the same ranking (Nigel: MLH of 0.231 and F of 0.271).

### DNA extraction, library preparation and genome sequencing

Multiple extraction methods were trialed to generate intact high-quality genomic DNA from stored muscle tissue of the single selected inbred shrimp. All DNA extractions and sequencing runs were carried out at the Australian Genome Research Facility (AGRF), Melbourne, Australia. For Illumina sequencing, the MagAttract HMW DNA kit (QIAGEN) was used and PCR-free fragment shotgun libraries were prepared using the ‘with-bead pond library’ construction protocol described by Fisher et al. (Fisher et al. 2011) with some modifications (Supplementary Material 1). The library was sequenced on two HiSeq 2500 lanes using a 250 bp PE Rapid sequencing kit (Illumina). The same DNA was also used to create a 10X Genomics Chromium library as per manufacturer instructions, which was sequenced on two HiSeq 2500 lanes using a 250 bp PE Rapid sequencing kit. For PacBio sequencing, the following DNA extraction methods were used with varying success: MagAttract HMW DNA kit (QIAGEN), Nanobind HMW Tissue DNA kit-alpha (Circulomics), and CTAB/Phenol/Chloroform (Supplementary Table 1). Libraries were prepared using the SMRTbell Template Prep Kit 1.0 (PacBio), loaded using either magbeads or diffusion, and sequenced using the Sequel Sequencing Kits versions 2.1 and 3.0 on a PacBio Sequel (Supplementary Table 1). The same muscle tissue was also used to prepare three Dovetail Hi-C libraries according to manufacturer’s instructions. Two libraries were sequenced on a shared lane of a NovaSeq S1 flow cell, and a third library was sequenced on one lane of a NovaSeq SP flow cell, with both sequencing runs generating 100 bp paired-end reads.

### Genome assembly

The quality of the initial short-read genome assemblies using either DISCOVAR *de novo* (Weisenfeld et al. 2014) with Illumina data, or Supernova (Weisenfeld et al. 2017) with 10X Genomics Chromium data was poor. The most contiguous assembly was achieved using wtdbg2/redbean (Version 2.4, Ruan and Li 2019) with 75 X times coverage of PacBio data, setting the estimated genome size to 2.2 Gb, but without using the wtdbg2 inbuilt polishing. The raw assembly was subjected to two rounds of polishing using the PacBio subreads data in arrow (Version 2.3.3, github.com/PacificBiosciences/GenomicConsensus) and one round of polishing using the Illumina short-read data in pilon (Version 1.23, Walker et al. 2014). Scaffolds were constructed in two steps. Medium-range scaffolding carried out using 10X Genomics Chromium data with longranger (Version 2.2.2, https://support.10xgenomics.com/genome-exome/software/downloads/latest) and ARCS (Version 1.0.6, Yeo et al. 2017), while long-range scaffolding was performed using dovetail Hi-C data, and intra- and inter-chromosomal contact maps were built using HiC-Pro (Version 2.11.1, Servant et al. 2015) and SALSA (commit version 974589f, Ghurye et al. 2017). This genome assembly was then submitted to NCBI GenBank, which required the removal of two small scaffolds and the splitting of one scaffold. The overall quality of the final V1.0 genome was assessed using BUSCO, and through mapping of RNA-seq, and Illumina short-reads using HiSAT2 (version 2.1.0, Kim et al. 2019).

### Repeat annotation

Repeat content was assessed with *de novo* searches using RepeatModeler (V2.0.1) and RepeatMasker (V4.1.0) via Dfam TE-Tools (V1.1, https://github.com/Dfam-consortium/TETools) within Singularity (V2.5.2, Kurtzer et al. 2017). Additionally, tandem repeat content was determined using Tandem Repeat Finder (V4.0.9, Benson 1999) within RepeatModeler. Analyses and plotting of interspersed repeats were carried out as per Cooke *et al.* (2020, github.com/iracooke/atenuis_wgs_pub/blob/master/09_repeats.md). Additionally, the genomes of the Black tiger shrimp (Thai origin, www.biotec.or.th/pmonodon; Kim et al. 2019), Whiteleg shrimp (*P. vannamei*, NCBI accession: QCYY00000000.1; Zhang et al. 2019), Japanese blue crab (*Portunus trituberculatus*, gigadb.org/dataset/100678; Tang et al. 2020), and Chinese mitten crab (*Eriocheir japonica sinensis*, NCBI accession: LQIF00000000.1) were run through the same analyses for comparison.

### Gene prediction and annotation

In order to generate an RNA-seq based transcriptome, raw data from a previous study (NCBI project PRJNA421400; Huerlimann et al. 2018) was mapped to the masked genome using STAR (Version 2.7.2b; Dobin et al. 2013), followed by Stringtie (Version 2.0.6; Pertea et al. 2015) (Supplementary Table 2). Additionally, the IsoSeq2 pipeline (PacBio) was used to process the ISO-seq data generated in this study (Supplementary Table 2). Finally, the genome annotation was carried out in MAKER2 (v2.31.10; Campbell et al. 2014; Cantarel et al. 2008; Holt and Yandell 2011) using the assembled RNA-seq and ISO-seq transcriptomes together with protein sequences of other arthropod species (Supplementary Table 3).

### Endogenous viral element analysis

BLASTn using a 3,832 bp IHHNV EVE Type A sequence detected in Australian *P. monodon* (Au2005; EU675312.1) as a query identified a potential EVE in Scaffold_97 of the *P. monodon* genome assembly. The EVE was unusual in that it comprised of repeated, inverted and jumbled fragments of an EVE Type A sequence. The nature and arrangement of EVE fragments was initially determined manually and the relative sequence positions of matching fragments within the EVE and scaffold sequence was determined using QIAGEN CLC Genomics Workbench 18.0 (https://digitalinsights.qiagen.com/). To confirm the authenticity of the Scaffold_97 EVE (S97-EVE), six PCR primer sets were designed using Primer 3 v.0.4.0 (Koressaar and Remm 2007; Untergasser et al. 2012) to amplify each EVE boundary and two internal sequences (Supplementary Table 4). DNA was extracted from ~10 mg gill tissue stored at −80°C from the *P. monodon* sequenced using DNAeasy kit spin columns (QIAGEN).

DNA was eluted in 50 μL EB buffer, aliquots were checked to DNA concentration and purity using a Nanodrop 8000 UV spectrophotometer and the remainder was stored at - 20°C. As DNA yields were low (9-38 ng/μL), a 1.0 μL aliquot of each sample was amplified in 10 μL reactions incubated at 30°C for 16 h as described in the REPLI-g Mini Kit (QIAGEN).

Each PCR (25 μL) contained 2 μL REPLI-g amplified gill DNA, 1 × MyTaq™ Red Mix (Bioline), 10 pmoles each primer and 0.25 μL (1.25 U) MyTaq DNA Polymerase (Bioline). Thermal cycling conditions were 95°C for 1 min followed by a 5-cycle touch-down (95°C for 30 s, 60°C to 56°C for 30 s, 72°C for 20 s), 30 cycles of the same using an anneal of 55°C for 30 s, followed by 72°C for 7 min and a 20°C hold. For semi-nested PCR using the 1b and 4b primer sets, 1 μL each PCR (either neat or diluted 1:5 to 1:10 depending on PCR product amount) was amplified similarly for 30 cycles using an anneal step of 55°C for 30 s. Aliquots (5-10 μL) of each reaction were electrophoresed in a 1.0% agarose-TAE gel containing 0.1 μL mL^−1^ ethidium bromide, and a gel image was captured using a Gel Doc 2000 UV transilluminator (Bio-Rad). Each amplicon was purified using a spin column (QIAGEN) and sequenced at the Australian Genome Research Facility (AGRF), Brisbane. The quality of sequence chromatograms was evaluated and consensus sequences for each amplicon were generated using Sequencher® 4.9 (Gene Codes Corp.).

### Data availability

Raw and assembled sequence data generated by this study have been deposited in GenBank BioProject PRJNA590309, BioSample SAMN13324362. PacBio and Illumina raw data can be found under accession numbers SRR10713990-SRR10714025. The final scaffolded assembly can be found under accession JAAFYK000000000. RNA-seq data used for annotation originated from an earlier study (Huerlimann et al. 2018).

## RESULTS AND DISCUSSION

### DNA extraction, library preparation and genome sequencing

In total, 158 Gb (72 X coverage) of Illumina, 494 Gb (224 X coverage) of 10X Genomics Chromium, 165 Gb (75 X coverage) of PacBio Sequel, and 119 Gb (54 X coverage) of DoveTail data were generated (Table 1). While the MagAttract HMW DNA kit (QIAGEN) was suitable for Illumina sequencing (PCR-free shotgun libraries and 10X Genomics Chromium), using this DNA resulted in poor PacBio Sequel sequencing runs (Supplementary Table 1). Runs consistently showed low yield and short fragment lengths, despite relatively high molecular weight DNA. However, DNA extracted with the Nanobind HMW Tissue DNA kit-alpha (Circulomics, Inc., Baltimore, USA) showed better sequencing performance (higher yield and fragment length; Supplementary Table 1). Furthermore, diffusion loading of the PB Sequel resulted in better results than magbead loading. DNA derived from either extraction method was unsuitable for Oxford Nanopore Technology (ONT) sequencing due to it rapidly blocking the pores (data not shown).

**Table 1.**
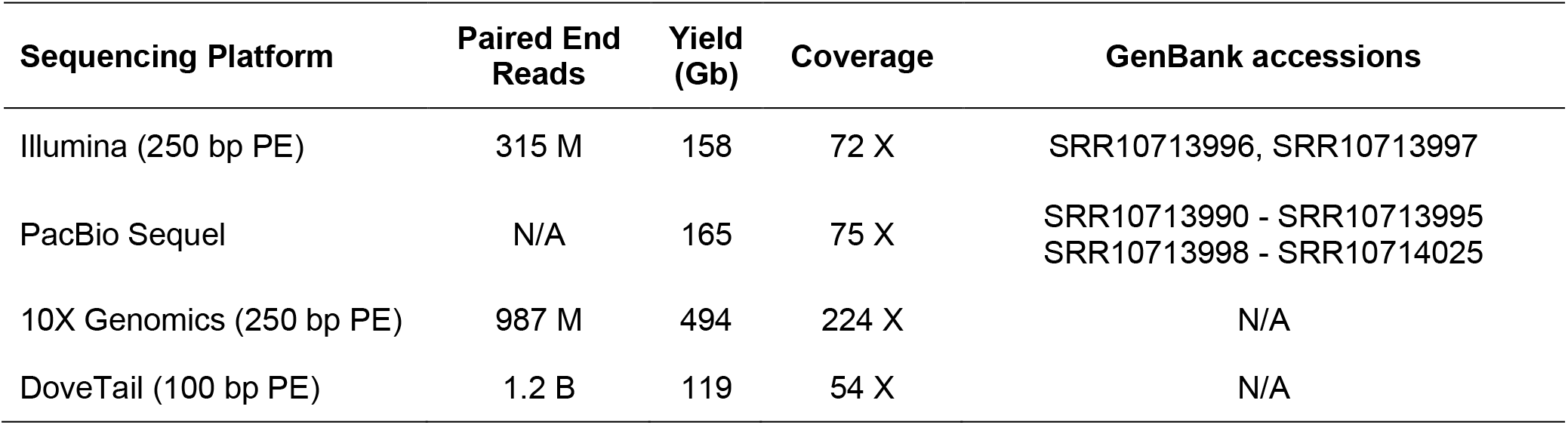
Illumina, PacBio, 10X Genomics, and DoveTail sequencing data used for the assembly and scaffolding of the black tiger shrimp genome.

Sequence quality issues associated with DNA extraction have also been noted in other shrimp genome assembly reports (Zhang et al. 2019; Uengwetwanit et al. 2020). The patterns seen in the PacBio sequencing results (short polymerase read lengths despite high quality libraries), coupled with the inability to successfully sequence *P. monod*on using ONT technology (immediate pore blockage), can be explained by high amounts of polysaccharides and polyphenolic proteins co-extracting with the DNA. This has also been mentioned by Angthong et al. (2020), who also present an alternative DNA extraction method to the Circulomics Nanobind HWM Tissue DNA extraction kit suggested here.

### Genome assembly and quality assessment

As reported by other Penaeid shrimp genome sequencing projects (Uengwetwanit et al. 2020; Zhang et al. 2019; Yuan et al. 2021a), sequencing and assembly of the Australian *P. monodon* genome proved problematic due to its large size, substantial heterozygosity and prevalence of repeat elements. The *de novo* assembly of the PacBio data resulted in 47,607 contigs (contig N50: 77,900 bp) a total of 1.90 Gbp in size (Table 2). After medium-range scaffolding with 10X Genomic Chromium data and long-range scaffolding with Dovetail sequences, the resulting scaffolded assembly contained 1.89 Gbp across 31,922 scaffolds (scaffold N50: *496*,398 bp; Table 2). Assuming a genome size of 2.2 Gbp (Huang et al. 2011), this scaffolded assembly covers 85.9 % of the projected *P. monodon* genome (Table 2). This is slightly lower than the 90.3% recently achieved for the same species in Thailand (Uengwetwanit et al. 2020), and higher than the 67.7% achieved for *P. vannamei* (Zhang et al. 2019), which has a slightly larger genome. Altogether, 98.1% of the Illumina DNA short-read data mapped to the raw assembly. BUSCO (V3; Simão et al. 2015), using the Arthropoda odb9 database (Zdobnov et al. 2017), estimated the Australian *P. monodon* genome assembly to be 86.8% complete (gene n = 1,066; 85.8% single copy; 1.0% duplicated; 4.5% fragmented; 8.7% missing; Table 2). These assembly metrics are comparable to those achieved for the Thai *P. monodon* assembly (C 87.9%, S 84.8%, D 3.1%, F 4.0%, M 8.0%; Uengwetwanit et al. 2020) and slightly better than those achieved for the *P. vannamei* assembly (C 78.0%, S 74.0%, D 4.0%, F 4.0%, M 18.0%; Zhang et al. 2019), both analyzed with the same database and BUSCO version (Table 2).

**Table 2.** Summary of assembly statistics for the Australian and Thai *P. monodon*, and *P. vannamei* genomes.

| Metrics | *P. monodon*<br>(Australia) | *P. monodon*<br>(Thailand) | *P. vannamei* |
| --- | --- | --- | --- |
| # contigs | 47,607 | 70,380 | 50,304 |
| Largest contig | 1,147,530 | 1,387,722 | 739,419 |
| Total length of contigs | 1.89 Gb | 2.39 Gb | 1.62 Gb |
| Contig N50 | 78 kb | 79 kb | 58 kb |
| # Scaffolds | 31,922 | 44 | - |
| Largest scaffold | 21.70 Mb | 65.87 Mb | - |
| Total length of scaffolds | 1.89 Gb | 1.99 Gb | 1.66 Gb |
| Scaffold N50 | 0.50 Mb | 49.0 Mb | 0.60 Mb |
| Projected Genome Size | 2.20 Gb | 2.20 Gb | 2.45 Gb |
| Percentage Covered By Scaffolds | 86.1% | 90.3% | 67.7% |
| GC (%) | 35.6 | 36.6 | 35.7 |
| Complete BUSCOs (C) | 86.8 | 87.9 | 78.0 |
| Complete and single-copy BUSCOs | 85.8 | 84.8 | 74.0 |
| Complete and duplicated BUSCOs | 1.0 | 3.1 | 4.0 |
| Fragmented BUSCOs (F) | 4.5 | 4.0 | 4.0 |
| Missing BUSCOs (M) | 8.7 | 8.0 | 18.0 |
| No. predicted gene models | 35,517 | 31,640 | 25,596 |
| No. of protein coding genes | 25,809 | 30,038 | - |
| No. genes annotated in | 17,158 | 20,615 | - |
| References | This study | Uengwetwanit et al. (2020) | Zhang et al. (2019) |

### Functional and repeat annotation

The functional annotation using RNA-seq, ISO-seq and protein information, identified 35,517 gene models, of which 25,809 were protein-coding and 17,158 were annotated using interproscan (Table 2). Similar numbers of genes were annotated in the Thai *P. monodon* (Uengwetwanit et al. 2020) and *P. vannamei* (Zhang et al. 2019) assemblies. Repeat content in the Australian *P. monodon* assembly (61.8%) was high, like in the Thai *P. monodon* assembly (62.5%; Uengwetwanit et al. 2020), and substantially higher than in genome assemblies of *P. vannamei* (51.7%; Zhang et al. 2019)), *Portunus trituberculatus* (45.9%, Tang et al. 2020) or *Eriocheir japonica sinensis* (35.5%, LQIF00000000.1) (Supplementary Table 5, Fig. 1). Interestingly, simple sequence repeats (SSRs) that dominated in prevalence (30.0%) in the Australian *P. monodon* assembly were less prevalent (23.9%) in the Thai *P. monodon* assembly (Uengwetwanit et al. 2020), similarly prevalent (27.1%) in the *P. vannamei* assembly, but far less prevalent in the genome assemblies of either the Japanese blue (16.9%) or Chinese mitten crab (7.9%) (Supplementary Table 5, Fig. 1). Such high SSR levels have been linked to genome plasticity and adaptive evolution facilitated through transposable elements (Yuan et al. 2021b). In addition to SSRs, the Australian *P. monodon* assembly contained 9.8% long interspersed nuclear elements (LINEs), 2.5% low complexity repeats, 2.0% DNA transposons, 1.6% long terminal repeats (LTRs), 0.51% small interspersed nuclear elements (SINEs), 0.1% satellites, 0.01% small RNA repeats and 15.4% unclassified repeat element types (Supplementary Table 5, Fig. 1). Broad comparisons of the major repeat types in the genome assemblies of *P. monodon*, *P. vannamei*, *Portunus trituberculatus* and *E. japonica sinensis* based on kimura distances showed them to be relatively conserved across all four crustacean species (Fig. 1). At all lengths and levels of divergence, unknown repeats dominated, with a large proportion of these >100 kbp in size (Fig. 1A). Repeat patterns shared across the four species were further highlighted when unknown reads were removed, and repeats split into major classes (Fig. 1B).

**Fig. 1.**
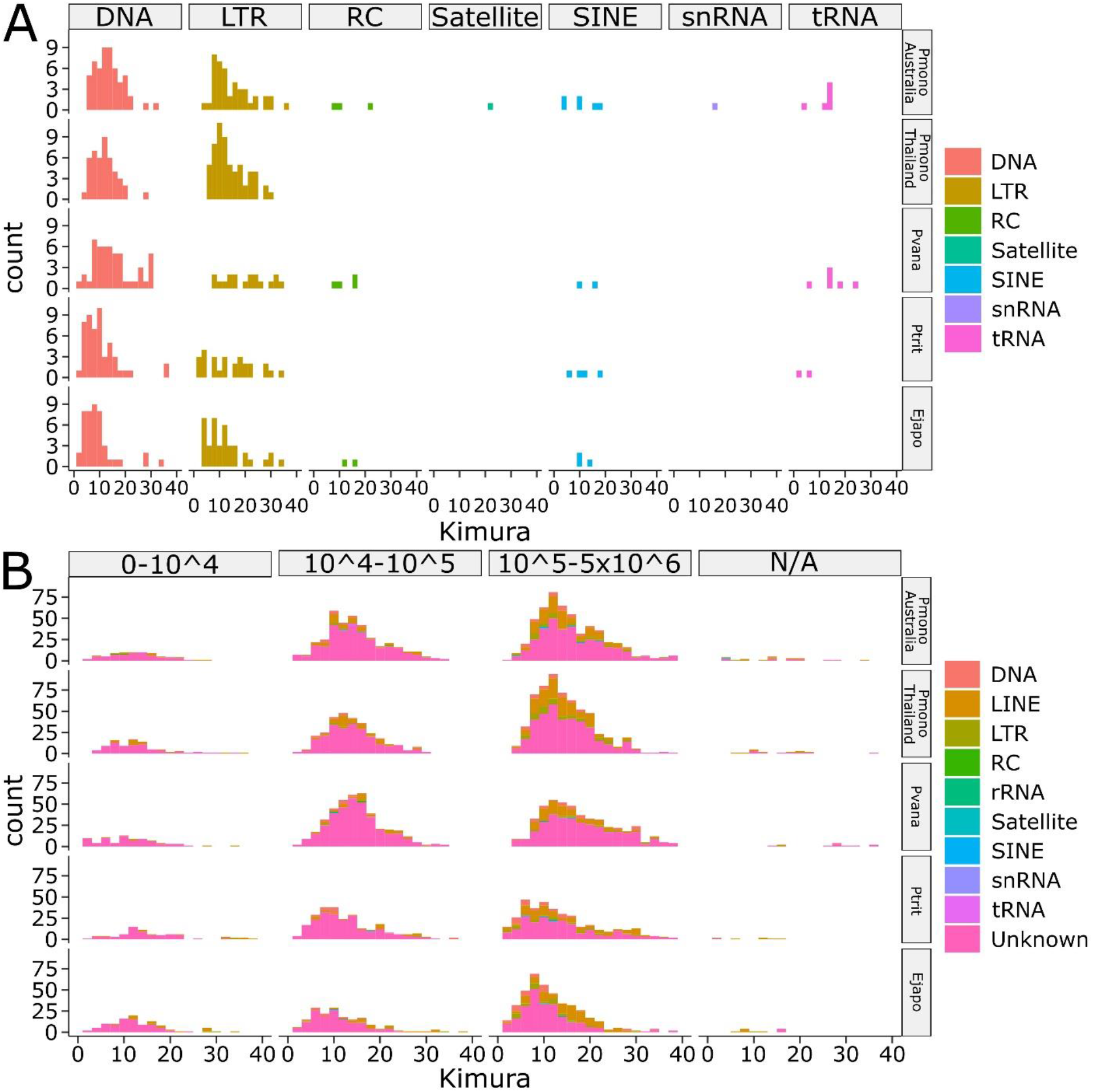
Kimura distances of repetitive sequences in the genome assemblies of Australian black tiger shrimp (*P. monodon*, NCBI accession: JAAFYK000000000, Pmono Australia, this study) Thai black tiger shrimp (Pmono Thailand, *P. monodon*, Pmono Thailand, Uengwetwanit et al. 2020), Whiteleg shrimp (Pvana, *Penaeus vannamei*, NCBI accession: QCYY00000000.1, Zhang et al. 2019), Japanese blue crab (Ptrit, *Portunus trituberculatus*, gigadb.org/dataset/100678, Tang et al. 2020), and Chinese mitten crab (Ejapo, *Eriocheir japonica sinensis*, NCBI accession: LQIF00000000.1) determined by using either (A) repeat length or (B) repeat class.

### IHHNV-EVE rearrangement in the Australian *P. monodon* genome

Sequences homologous to a 3,832 bp linear IHHNV-EVE (Au2005, Type A) found to occur in some Australian *P. monodon* (Krabsetsve et al. 2004) were identified in Scaffold_97 (S97, 2,608,951 nt). However, rather than representing an intact linear copy of this EVE, the S97-EVE comprised a 9,045 bp stretch of jumbled, repeated, and inverted IHHNV fragments flanked by two repeated 591/590 bp (flanking repeat) sequences (Fig. 2). Alignments identified most fragments to be jumbled relative to their location in the Au2005 IHHNV-EVE sequence, and the expanded EVE length to be due to replicated short sequences originating from 5’-terminal genome regions. Fragments positioned at the S97-EVE extremities generally originated from the central and downstream regions of the Au2005 IHHNV-EVE sequence and were consistently orientated inwards. The central S97-EVE region comprised a block of at least six 661 bp repeat units (RUs). Each RU was comprised of two inward-facing sequences either (A) 398 bp or (B) 263 bp in length that mapped to the same region (94-501 and 94-368, respectively) at the 5’-terminus of the Au2005 IHHNV-EVE (Fig. 2B, grey arrows). In total, 83% of the Au2005 IHHNV-EVE sequence was identified to be covered by genome fragments present in the S97-EVE, with those present being on average 99.3% identical.

**Fig. 2.**
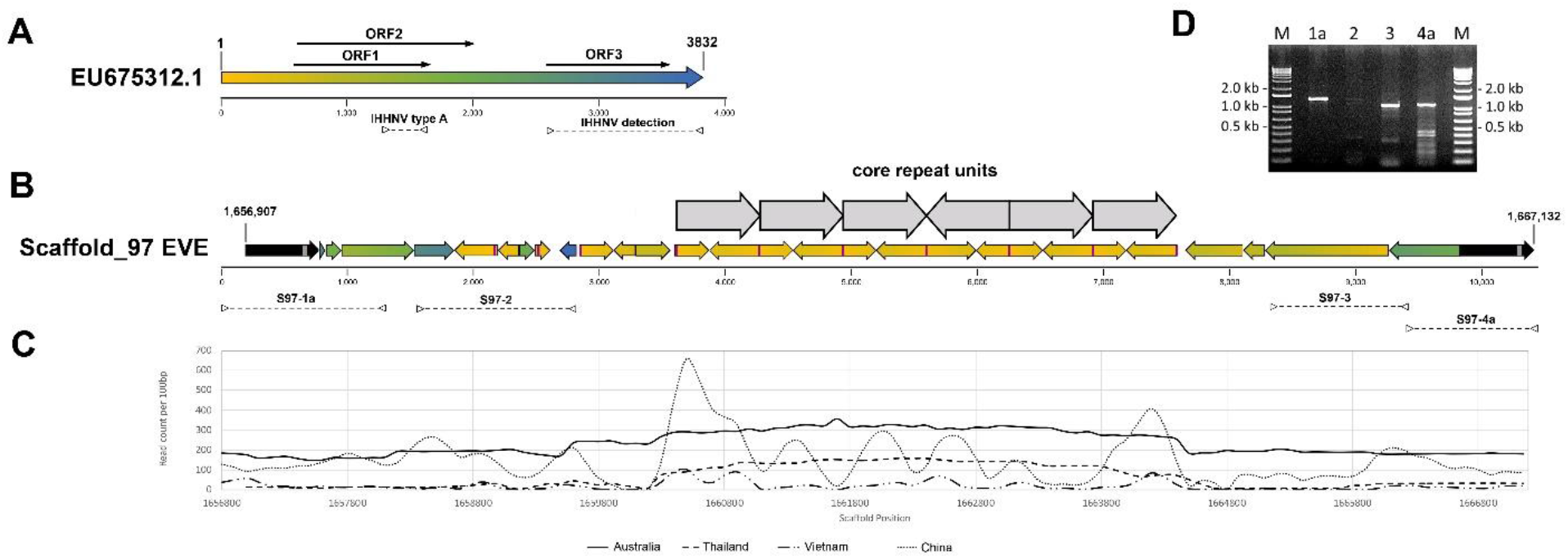
(A) Schematic diagram of a 3,832 bp ssDNA genome of Infectious hypodermal and hematopoietic necrosis virus (IHHNV) showing the relative positions of coding sequences (arrows) for the virus replicase (ORF1), NS1 non-structural protein (ORF2) and viral capsid protein (ORF3). A colour gradient was applied to visualize relative genome positions. (B) Schematic diagram of the positions and orientations of IHHNV genome fragments comprising the Scaffold_97 EVE (S97-EVE). The orientations of the IHHNV fragments (coloured arrows) and the flanking repeated 591/590 bp host sequence (black arrows) are shown by arrow directions. The origins of the S97-EVE fragments relative to their positions in a linear IHHNV-EVE (see A) are identified by colour. The 10,226 bp S97-EVE resided between positions 1,656,907 and 1,667,132 in the 2,608,951 bp Scaffold_97 sequence. The larger grey arrows identify the positions and orientations of at least 6 core repeat blocks comprising of 2 smaller inverted repeats. Grey vertical bars show the location of a 34 bp sequence in each flanking repeat capable of folding into a stable secondary structure. The purple vertical bars show the locations of the 18 bp palindromic sequence present at the boundaries of each repeat unit (RU) and partial RU. Dashed lines (>--<) identify the regions amplified by the 4 PCR tests S97-1a, S97-2, S97-3, and S97-4a. (C) Coverage depth across the S97-EVE sequence of raw short reads used to assemble genome scaffolds of *P. monodon* from Australia (this study), Thailand (Uengwetwanit et al. 2020), Vietnam (Van Quyen et al. 2020) and China (Yuan et al. 2018). (D) Agarose gel image showing DNA products amplified by the S97-1a, S97-2, S97-3, and S97-4a PCR tests.

The inverted A and B sequences comprising each RU contain RNA transcription regulatory signals of the IHHNV P2 promoter (Shike et al. 2000; Dhar et al. 2011; Dhar et al. 2010; Dhar et al. 2007). Both initiated at a sequence (5’-GTCATAGGT…) mapping precisely to a G nucleotide residing immediately downstream of the inversion point (|) of an 18 bp <u>inverted repeat</u> (5’-..TTACA<u>ACCTATGAC|GTCATAGGT</u>CCTATATAAGAGT..-3’) located 2 bp upstream of the TATA-box element (5’-TATATAA-3’) of the P2 transcriptional promoter (Dhar et al. 2011; Dhar et al. 2010; Dhar et al. 2007). The A and B repeat components in each RU of the six blocks were orientated 5’|B-A|B-A|B-A|A-B|B-A|B-A|3’, with those in RU4 being reversed compared to the others. Due to the A and B repeat components being inverted, the 18 bp inverted repeat (ie. 5’-..ACTCTTATATAGG<u>ACCTATGAC|GTCATAGGT</u>CCTATATAAGAGT..-3’) was reconstructed at each of the 5 RU junction sites irrespective of which 2 repeat components (A|A, A|B or B|B) were joined (Fig. 2B, purple bars). This arrangement generated a 544 bp inverted repeat (263 × 2 + 18) for sequences extending from either A|B or B|B RU junctions, or a 1,902 bp inverted repeat (661 × 2 + 263 × 2 + 18 × 3) for the long complimentary sequence stretches extending outwards from the A|A components at the RU3|RU4 junction to the end of repeat component A of RU2 and the equivalent position of repeat component B in RU5. However, relating to the descriptions of this unusual EVE segment, it is important to note that no single long read was obtained that traversed the entire six RU blocks into flanking unique S97-EVE sequences (Fig. 2). Combined with short read numbers generated using various sequencing methods being substantially elevated at positions mapping to each block RU (Fig. 2C), the likelihood of the block comprising more than six RUs remains to be established.

DNAFold and RNAfold analyses showed the 18 bp inverted repeat, the inverted A and B repeat components of each RU and the longer complimentary sequences that stretched through multiple RUs to all have potential to form highly stable simple to complex secondary structures as either ssDNA or ssRNA (data not shown). Discrete DNA secondary structures are known to have roles in mediating recombination in mobile genetic elements (Bikard et al. 2010) and in the genomes of parvoviruses like the extensively studied adeno-associated virus (AAV), structures formed by inverted terminal repeat (ITR) sequences play critical roles in initiating genomic ssDNA replication, genomes forming circular extrachromosomal dsDNA episomes and genomic integrating into host chromosomal DNA (Cotmore and Tattersall 1996, Kotin et al. 1991, Schnepp et al. 2005, Yang et al. 1997). The mechanisms leading to the A and B inverted repeat sequences forming the 661 bp RUs and their apparent multiplication in the central region of the S97-EVE remains unknown. However, their existence is consistent with integrated AAV proviral DNA structures being observed to contain head-to-tail tandem arrays of partial ITR sequences and for genomic rearrangements occurring via deletion and/or rearrangement-translocation at the integration site (Yang et al. 1997).

The 18 bp inverted repeat at the S97-EVE RU junctions also occurred at the upstream RU1 and downstream RU6 boundaries of the RU block. However, unlike those at the internal RU junctions which extended into the same downstream Au2005-EVE sequence including the TATA-box element (Dhar et al. 2011; Dhar et al. 2010; Dhar et al. 2007; Krabsetsve et al. 2004), the outer half of each inverted repeat flanking the RU-block extended into sequences toward the 5’ end of the IHHNV genome (Supplementary Figure 1). Three disparate partial RU sequences (pRUa, pRUb, pRUc) associated with four 18 bp inverted repeats also resided just upstream of the 6 RU block. Like RU1 and RU6, one side of each inverted repeat possessed variable lengths of sequence extending toward the IHHNV genome 5’-terminus (Supplementary Figure 1).

In some IHHNV strains, the sequence immediately upstream of the 18 bp inverted repeat comprises a second imperfect 39-40 bp inverted repeat. With an IHHNV strain detected in Pacific blue shrimp (*Penaeus stylirostris*) sampled from the Gulf of California in 1998 (Shike et al. 2000, AF273215.1), the 5’-genome terminus upstream of it consisted of an 8 bp portion of the downstream 18 bp inverted repeat (Supplementary Figure 1). In the S97-EVE, the 18 bp inverted repeats associated with each terminal RU or upstream pRU extended 18-38 bp into the 39-40 bp inverted repeat (Supplementary Figure 1). Of interest, with the first pRU occurring in the S97-EVE (5’-pRUa), the 93 bp sequence abutting the 18 bp inverted repeat was also identical to the 5’-terminal sequence reported for the Au2005 IHHNV-EVE found in *P. monodon* sampled from farms in Australia in 1993/1997 (Krabsetsve et al. 2004; EU675312.1).

To confirm that the fragmented and jumbled nature of the S97-EVE was not an assembly artefact, regions spanning each EVE extremity to unique host sequences positioned just beyond the 591/590 bp flanking repeats, as well as two internal regions each spanning conjoined non-repeated EVE fragments were amplified by PCR (Supplementary Table 4; Fig. 2D). Amplicons of the expected sizes were clearly amplified by each extremity PCR test (S97-1a and S97-4a) and the S97-3 internal PCR test (Fig. 2D). The other internal PCR test (S97-2) also generated a 1,337 bp amplicon of the expected size, as well as one ~200 bp shorter, but in relatively lower abundance. Using each extremity PCR product as template, semi-nested PCR tests using an alternative internal EVE-specific primer also produced amplicons of the expected shorter sizes, and their authenticity was confirmed by sequence analysis (data not shown).

### *P. monodon* repeat sequences flanking the IHHNV-EVE

BLASTn and BLASTx searches did not identify any homologues of the 591/590 bp flanking repeat sequence in GenBank. However, searches of the *P. monodon* genome assembly identified long closely-related sequences in hundreds of other scaffolds (data not shown). The searches also highlighted the presence of a 34 bp sequence (5’-..<u>ATGACTCC</u>T<u>CCCCC</u>ATAGATA<u>GGGG</u>C<u>GGAGTCAT</u>..-3’) in each flanking repeat (Fig. 2B, grey bars, upstream repeat position 1,657,364-1,657,397; downstream repeat position 1,667,000-1,667,033) that was also present in 178 other scaffolds at >80% identity. DNAFold and RNAfold analyses showed the sequence and its reverse compliment to fold into stable hairpin structures as either ssDNA (ΔG = −10.44/−11.92, Tm = 83.8/85.7°C) or ssRNA (ΔG = −20.40/−23.70). However, whether this or other sequences in the host flanking repeat interact with IHHNV genome sequences and proteins to facilitate recombination and site-specific integration remains to be investigated. In this regard, the flanking host repeat possessed a 5’..<u>CTTACTTA</u>CA<u>CTTG</u>..3’ tetramer repeat, which to the 5’-side of the S97-EVE was located 33 bp upstream of the IHHNV <u>CTTA</u>.. sequence at the host/S97-EVE junction, much like the host tetramer repeats well characterised to be pivotal to the AAV genome integrating at a specific location in human chromosome 19 (Kotin et al. 1992, Linden et al. 1996).

### Comparison to jumbled IHHNV-EVEs in other *P. monodon* genome assemblies

BLASTn searches of the most comprehensive genome assembly of a *P. monodon* from Thailand (NSTDA_Pmon_1, GCA_015228065.1, Uengwetwanit et al. 2020) identified Scaffold_35 (S35) containing two disparate aggregations of jumbled IHHNV-EVE Type A fragments (S35-EVE1 = 7,888 bp; S35-EVE2 = 16,310 bp) each flanked by >500 bp host repeats near identical in sequence to those flanking the S97-EVE (Table 3). Compared to the S97-EVE, 2,328 bp of S35-EVE1 sequence immediately downstream of the 5’ 592 bp host repeat, except for a 166 bp deletion, and 647 bp of sequence immediately upstream of the 3’ 591 bp host repeat, were identical. Further inwards, however, the order and arrangement of EVE fragments diverged.

**Table 3.** Detection and notable features of IHHNV-EVE sequences identified in other genomes of *P. monodon*.

| Reference Genome IDs | Notable EVE features |  |  |  |  |
| --- | --- | --- | --- | --- | --- |
|  | Start | End | Length (bp) | Orientation | Homology (%) |
| *P. monodon* Thailand (Uengwetwanit et al. 2020) |  |  |  |  |  |
| *Scaffold 35 EVE-1* | 770,236 | 778,124 | 7,888 |  |  |
| RU1 | 772,730 | 773,391 | 661 | minus | 99.9 |
| RU2 | 773,450 | 774,111 | 661 | plus | 100.0 |
| RU3 | 774,170 | 774,831 | 661 | minus | 99.9 |
| RU4 | 774,890 | 775,551 | 661 | plus | 97.9 |
| *Scaffold 35 EVE-2* | 862,618 | 878,928 | 16,310 |  |  |
| RU1 | 866,534 | 867,145 | 611 | minus | 79.4 |
| RU2 | 867,204 | 867,791 | 587 | plus | 81.3 |
| RU3 | 867,840 | 868,467 | 627 | minus | 83.5 |
| RU4 | 868,515 | 869,130 | 615 | plus | 80.0 |
| RU5 | 872,127 | 872,754 | 627 | plus | 78.9 |
| RU6 | 872,799 | 873,434 | 635 | minus | 90.0 |
| RU7 | 873,492 | 874,152 | 660 | plus | 97.2 |
| RU8 | 875,469 | 876,168 | 699 | plus | 92.1 |
| *P. monodon* Vietnam (Pmod26D_v1; GCA_007890405.1) |  |  |  |  |  |
| VIGR010059916.1 EVE (4,003 bp) |  |  | 4,003 |  | 98.4 |
| VIGR010211091.1 EVE (1,917 bp) |  |  | 1,917 |  | 99.0 |
| VIGR010168684.1 EVE (2,220 bp) |  |  | 2,220 |  | 98.9 |
| *P. monodon* China (Pmon_WGS_v1, GCA_002291185.1) |  |  |  |  |  |
| gb\|NIUS011382605.1 (645 bp) |  |  | 645 |  | 98.9 |
| gb\|NIUS011109800.1 (848 bp) |  |  | 848 |  | 98.3 |

As in the S97-EVE, the central region of the S35-EVE1 contained a block of 4 × 661 bp RUs each comprised of the same inward facing (A) 398 bp and (B) 263 bp repeats but ordered 5’|A-B|B-A|A-B|B-A|3’, thus making a 2877 bp inverted repeat with an inversion point at the RU2-RU3 boundary. Also, like the 97-EVE, each S35-EVE RU was flanked by same 18 bp inverted repeat sequence, with those upstream of RU1 and downstream of RU4 extending 17-33 bp into a 41 bp imperfect inverted repeat sequence located immediately upstream toward the 5’-genome termini in some IHHNV strains (Supplementary Figure 2). However, unlike the RU block in the S97-EVE, each of the three internal S35-EVE RU boundaries comprised of 2 × 18 bp inverted repeats flanking the complete 41 bp imperfect inverted repeat (Supplementary Figure 2). This revised the RU junction to the inversion point in longer imperfect inverted repeat, rather than the inversion point of the 18 bp inverted repeat. DNAfold and RNAfold analyses showed that the 41 bp inverted repeat and its reverse compliment sequence could fold into stable hairpin structures as either ssDNA (ΔG = −14.18/−14.86, Tm = 73.6/75.6°C) or ssRNA (ΔG = −22.50/−25.00).

The larger S35-EVE2 sequence differed in the arrangement and homology of up to eight RUs, possibly composed of two entirely duplicated inward-facing EVE fragments (Table 3). The IHHNV-EVE fragments in S35-EVE1 contained 72% of the Au2005 IHHNV-EVE sequence with 98.8% homology, on average. In contrast, IHHNV-EVE fragments in S35-EVE2 region only contained 53% of the IHHNV-EVE sequence with 97.5% homology, on average.

BLASTn searches of the genome assembly of a *P. monodon* from Vietnam (Pmod26D_v1, GCA_007890405.1, Van Quyen et al. 2020), using the 9,045 bp S97-EVE and 3,832 bp linear Au2005 Type A IHHNV-EVE sequences identified 3 short contigs (*VIGR010059916.1, 4,003 nt; VIGR010168684.1, 2,220 nt; VIGR010211091.1,* 1,917 bp) also comprised of jumbled IHHNV-EVE Type A-like fragments (Table 3). In two of the contigs, the stretches of jumbled EVE fragments neighbored either a complete (590 bp) or incomplete (356 bp) host repeat sequences like those flanking the S97-EVE. BLASTn searches of a genome assembly of a *P. monodon* from Shenzhen, China (Pmon_WGS_v1, GCA_002291185.1) also identified evidence of an EVE comprised of jumbled IHHNV genome fragments (Table 3), and despite contig lengths being short, it was also being flanked by the same repeated host sequence flanking the S97-EVE (data not shown). While more complete higher quality genome assemblies would add confidence, the insertion locations of the jumbled EVEs present in the genome assemblies of the *P. monodon* from Vietnam and China appear shared with those the Australian S97-EVE and Thai S35-EVE1, with the second less-related jumbled S35-EVE2 in the Thai genome residing at a nearby site. Interestingly, BLASTn searches of the genome assemblies of *P. monodon* from Australia, Thailand, Vietnam, or China identified no evidence of linear IHHNV-EVE forms.

### Origins and implications of jumbled IHHNV-EVEs

While varying in lengths, the amalgamations of reordered, inverted, and repeated IHHNV genome fragments comprising the EVEs detected in Scaffold_97 (S97) of the Australian *P. monodon* assembly (this study) and in Scaffold_35 (S35) of the Thai *P. monodon* assembly (Uengwetwanit et al. 2020) share an integration site as well as structural and sequence similarities with the partial EVE sequences detected in short contigs of genome assemblies of *P. monodon* originating from Vietnam and China (as outlined above). These similarities are suggestive of a progenitor IHHNV genome becoming stably integrated as an EVE prior to *P. monodon* becoming dispersed widely across its current distribution range. Such an ancient event would also support differences noted for example in EVE fragment composition, central RU numbers, and the nature of the conserved inverted-repeat sequences defining the boundaries of the RUs. Furthermore, the conservation of the inverted-repeat sequences at the RU boundaries and their potential to form stable ssDNA folding structures suggests a potential role in their apparent multiplication.

The IHHNV P2 RNA transcriptional promoter motifs, including the 18 bp inverted repeat sequences and TATA-box (Dhar et al. 2014; Shike et al. 2000; Silva et al. 2014), at the RU boundaries have potential to facilitate transcription of various virus-specific sense and antisense ssRNA sequences. RNA transcribed from them would then be capable of forming long virus-specific dsRNA or hairpin dsRNAs, potentially in high abundance due to their repeated nature. If so, such virus-specific antisense RNAs or dsRNA forms processed through the RNA interference (RNAi) machinery of *P. monodon* (Attasart et al. 2010; Attasart et al. 2011; Dhar et al. 2014; Su et al. 2008) could provide resilience against IHHNV infections progressing to become acute and cause disease. Such an advantage might promote the selection of *P. monodon* carrying this form of IHHNV-EVE, particularly in circumstances when shrimp are specifically selected or bred for aquaculture robustness. Selection for the EVE over several years would also be consistent with the viral accommodation model hypothesized to involve farmed shrimp acquiring and/or selected for an ability to mount elevated antisense ssRNA-based and/or dsRNA-based anti-viral responses (Flegel 2007, 2020; Flegel 2009).

EVEs comprised of reordered, inverted, repeated and missing IHHNV genome fragments would be expected to invalidate many PCR tests either designed specifically, or found through use, to amplify IHHNV-EVE dsDNA sequences (Cowley et al. 2018; Rai et al. 2009; Rai et al. 2012; Saksmerprome et al. 2011; Tang et al. 2007). As examples, the 356 bp sequence targeted by the 77102F/77353R primer set (Nunan et al. 2000) found to amplify both viral ssDNA and EVE dsDNA sequences existed in the S97-EVE and S35-EVE1, but not in the S35-EVE2 sequence. However, nucleotide mismatches at the 3’ terminal position of both primers and at four other positions in the 18-mer 77353R primer would likely compromise the capacity of this primer set test to amplify these EVEs. In contrast, neither EVE sequence possessed intact fragments spanning regions amplified by primer sets 392F/R (392 bp) and 389F/R (389 bp) recommended by the OIE as useful for amplifying divergent IHHNV strains as well as IHHNV-EVE Type A and B sequences, or primer set MG831F/R (831 bp) designed specifically to amplify known linear IHHNV-EVE types (Tang et al. 2007). Similarly, the region targeted by a real-time PCR primer set designed to specifically amplify IHHNV-EVE Type A sequences was absent from the S97-EVE and S35-EVE1, but present, albeit with some primer mismatches, in the S35-EVE2 sequence (Cowley et al., 2018).

Variability among individual *P. monodon* in EVE sequences amplified by a suite of 10 PCR primer sets covering overlapping regions of complete linear IHHNV-EVE sequence have been interpreted to suggest the random integration of IHHNV genome fragments (Saksmerprome et al. 2011). While the jumbled fragments in the IHHNV-EVEs described here might explain these, the diversity in EVE makeup suggested by these data would require jumbled EVEs to be characterized in larger numbers of *P. monodon,* or other penaeid species susceptible to IHHNV infection. Such broader information will also be important to devising PCR methods to detect jumbled IHHNV-EVE sequences more reliably.

## Conclusions

Using PacBio long-read data with Illumina short-read polishing together with 10X Genomics and Hi-C scaffolding, this study generated a draft genome assembly and annotation of a black tiger shrimp (*Penaeus monodon*) originating from Australia. The assembly represents the first to be produced from this geographically isolated and genetically distinct population (Vu et al. 2021). The assembly therefore adds to the genetic resources available for *P. monodon* and Penaeid shrimp in general, and will assist investigations into their evolution and genome expansion resulting from transposable elements. Of the *P. monodon* genome features, the high prevalence of general repeats is the most remarkable, and especially the high content of SSRs even in comparison to other crustacean species. Another unexpected feature was the existence of a previously undescribed IHHNV endogenous viral element (EVE) located between a repeated host sequence. Rather than being comprising of a linear sequence of all or part of the ~3.9 kb IHHNV genome, the EVE comprised of a conglomerate of reordered, inverted, and repeated IHHNV genome fragments. Searches of genome assemblies available for *P. monodon* from Thailand, Vietnam and China indicated with variable confidence, depending on assembly quality, that each contained a similarly jumbled IHHNV-EVE inserted at the same genome location. The fragmented and rearranged nature of these EVEs has implications for detecting them with currently available PCR tests. The presence of multiple inverted sequences including multiple IHHNV RNA transcription promoter elements also has implications for them expressing virus-specific dsRNA capable of interfering with exogenous IHHNV replication. The complexity of the rearranged IHHNV genome fragments comprising the EVEs begs many questions related to how long they have existed in the genomes of genetically diverse *P. monodon*, as well as to what processes have led to their integration at a specific genome location, to the IHHNV genome fragments becoming rearranged and to the apparent multiplication of a repeat unit comprised of highly defined inverted sequences derived from the 5’-terminal region of the IHHNV genome.

## Acknowledgments

This research would not have been possible without the significate in-kind contribution and support by Seafarms Group Ltd, including the acquisition of the G1 animals, carrying out the breeding and providing the necessary facilities and expertise. The authors also thank Min Rao for technical assistance in generating and sequencing S97-EVE PCR amplicons.

## Funding

This work was funded by the Australian Research Council (ARC) Industrial Transformation Research Hub scheme, awarded to James Cook University (Grant ID: IH130200013) and in collaboration with the Commonwealth Scientific Industrial Research Organisation (CSIRO), the Australian Genome Research Facility (AGRF), the University of Sydney and Seafarms Group Pty Ltd. AGRF is supported by the Australian Government National Collaborative Research Infrastructure Initiative through Bioplatforms Australia”.

